# Targeting cardiac myocyte Na^+^-K^+^ pump function with β3 adrenergic agonist in rabbits and patients with severe congestive heart failure

**DOI:** 10.1101/804245

**Authors:** Natasha AS Fry, Chia-Chi Liu, Alvaro Garcia, Elisha J Hamilton, Keyvan Karimi Galougahi, Yeon Jae Kim, Jacqueline L Harris, Gregory IC Nelson, David W Whalley, Henning Bundgaard, Helge H Rasmussen

## Abstract

**Background:** Reported one- and two-year mortality for patients with advanced heart failure (HF) treated medically are ~75% and nearly 100%. In such patients, reversible cellular abnormalities are potential treatment targets and a raised cytosolic Na^+^ concentration that impairs their myocardial contractility is one potential target. β3 adrenoceptor (β3 AR) agonists stimulate the myocyte Na^+^-K^+^ pump.

**Methods:** We induced severe HF in rabbits by coronary ligation and measured indices of organ congestion after treatment with β3 AR agonists. Na^+^-K^+^ pump current was measured in voltage-clamped myocytes isolated from non-infarct myocardium. To assess if β3 ARs might add benefit to optimised guideline-directed medical treatment we report outcomes of giving the β3 AR agonist mirabegron to patients hospitalized with advanced, treatment-refractory stage D HF.

**Results:** Treatment of rabbits after coronary ligation with β3 AR agonist reversed a decreased myocyte Na^+^-K^+^ pump current and significantly reduced organ congestion and prevalence of ascites. Oral treatment with mirabegron rapidly improved signs and symptoms of 9 patients with advanced HF and improvement of ≥1 NYHA Class was maintained early post-discharge with continued treatment. One patient died from HF at 16 months, 4 died from other causes at 2 – 30 months and 4 remain alive at 38 ± 4 months with NYHA Class II symptoms.

**Conclusions:** Parallel β3 AR agonist-induced reversal of Na^+^-K^+^ pump inhibition and severe HF in rabbits identify pump inhibition as a treatment target, and changed in-hospital clinical trajectory and post-discharge course more favorable than expected suggest efficacy of mirabegron in advanced human HF.

## Introduction

Even in end-stage heart failure (HF), cellular abnormalities in viable but dysfunctional myocardium are hypothesized to be useful treatment targets ^1, 2^ and contractile dysfunction of myocardium isolated from hearts of patients with end-stage HF with reduced ejection fraction (HFrEF) is at least in part due to a pathologically high cytosolic Na^+^ concentration ([Na^+^_i_]) ^3^. A decrease in the raised [Na^+^_i_] can be achieved by activation of its key determinant, the Na^+^-K^+^ pump.

The cardiac myocyte Na^+^-K^+^ pump is activated in vitro by agonists for the β3 adrenoceptor (β3 AR) ^4^, and an agonist given acutely by intravenous infusion to 4 normal sheep decreased cardiac contractility as indicated by hemodynamic indices. After severe HFrEF was experimentally induced, the agonist shifted indices towards an increase in contractility ^4^. While such differential effects in normal vs. failing hearts are consistent with β3 AR agonist-induced the Na^+^-K^+^ pump activation and a decrease in [Na^+^_i_] ^3^, raised neurohormone levels in HF are expected to cause Na^+^-K^+^ pump inhibition ^5^. It cannot be assumed β3 AR agonist can counter this in vivo.

Here we have directly measured effects of in vivo β3 AR agonist treatment on myocyte Na^+^-K^+^ pump function in rabbits with HFrEF. While hemodynamic indices in the sheep model suggested a β3 AR agonist was beneficial in HFrEF such indices are poor predictors of treatment outcomes in HF ^6^. To address more relevant clinical objectives we studied effects of treatment with β3 AR agonists on congestive indices in the rabbits. Myocyte Na^+^-K^+^ pump function was reduced in rabbits with HF. Pump inhibition was eliminated by β3 AR agonist treatment and, in parallel, congestive indices improved.

The rabbits had severe HF as indicated by marked organ congestion and ascites, but structural and functional remodelling of long-standing human HF are not necessarily reproduced in animal models. In addition, it cannot be established if the rabbit HF was equivalent to the *end-stage* human HF in which a high [Na^+^_i_] caused contractile myocardial dysfunction ^3^. Also, efficacy of monotherapy in the rabbits does not imply added efficacy to optimized guideline-directed treatment. No animal model can reproduce this clinical trial practice or the setting of co-morbidities often limiting treatment options in advanced human HF.

We complement our studies on the rabbits by reporting on outcomes of adding the β3 AR agonist mirabegron to treatment of hospitalized patients with advanced, stage D HF. There is a paucity of studies describing outcomes for such patients ^7^. However, one reported that of medically treated patients, not undergoing cardiac transplantation or having mechanical circulatory support (MCS), 75% had died by 1 year and almost all by 2 years, more than 90% from heart failure ^8^.

Our rationale for prescribing mirabegron was in part based on our previous study on patients with stable, mostly Class II NYHA HFrEF. Mirabegron seemed safe, and an exploratory secondary analysis suggested treatment increased left ventricular ejection fraction (LVEF) in patients with a particularly low baseline EF, but not in similar placebo treated patients ^9^. There is also a plausible rationale for efficacy of β3 AR agonists in advanced HF. Most conventional drugs used in treating HFrEF prevent Na^+^–K^+^ pump inhibition that neurohormonal activation causes ^5^ and they are often not tolerated in target doses or not tolerated at all in advanced HF. Rather than preventing Na^+^–K^+^ pump inhibition β3 AR agonists directly stimulate the pump ^4^, that is, they address pathophysiology that remains after conventional treatment. In considering treating patients with mirabegron we also took into account that there is no effective medical treatment for advanced HF ^7, 10^.

Mirabegron is approved for treatment of overactive bladder and as such was prescribed off-label. At least two cardiologists had to agree the HF was refractory to maximal tolerated guideline-directed medical treatment and that it was unlikely that the patients could be discharged from hospital. We report in-hospital clinical trajectories before and after we started treatment with mirabegron and post-discharge outcomes during maintained treatment.

## Methods

Rabbits were used because β3 AR agonist affinities in rabbit cardiac myocytes and functional effects of activation of it are similar to those in human myocardium ^11^. We induced HFrEF by ligating the circumflex coronary artery. Details of anaesthesia and postoperative analgesia and experimental protocols are in Expanded Methods in the Data Supplement. Protocols were designed to minimise the number of rabbits and were approved by the Royal North Shore Hospital Animal Ethics Committee.

Echocardiography was performed before coronary artery ligation or sham operations and 7 days after surgery. A LVEF <25% was used to decide if rabbits should enter the treatment protocols because we had found a value <25% to be a reliable marker of HF at subsequent autopsy as indicated by organ congestion and an increase in LV mRNA levels for brain natriuretic peptide (BNP).

We used pulmonary congestion as reflected by lung: body weight ratio as the primary endpoint. This is supported by a strong correlation between lung weight at autopsy and extravascular lung water in vivo, robust across diverse co-morbidities in humans ^12^. We also recorded liver and heart weights and presence or absence of ascites. LV BNP mRNA was not used as an outcome variable because we expected it to change too slowly for detection of changes within treatment periods of 3 or 14 days that were used. Furthermore, serum BNP in clinical studies are poor surrogates for treatment efficacy in decompensated HF ^6^.

We started treatment with the β3 AR agonists CL316,243 infused via subcutaneous osmotic minipumps at 2 mg/kg/24 hours ≥2 weeks after coronary ligation. CL316,243 is a selective agonist for the rabbit β3 AR ^11^ and it stimulates the Na^+^-K^+^ pump in isolated rabbit cardiac myocytes ^4^. Duration of treatment was limited to 3 days due to prohibitive cost of the drug in a medium sized animal like rabbit. We treated another group of rabbits for 2 weeks with the β3 AR agonist ASP9531 (a gift from Astellas Pharma) via osmotic minipumps at a rate of 250 μg/kg/day.

Electrogenic Na^+^-K^+^ pump current (I_p_, arising from the 3: 2 Na^+^: K^+^ exchange ratio) was measured in myocytes from non-infarcted myocardium using the whole-cell patch clamp technique. Patch pipette solutions perfusing the intracellular compartment included 10 mmol/L Na^+^. I_p_ was identified according to pre-determined quality criteria ^13^. I_p_ (in pA) is reported normalized to membrane capacitance (in pF) and hence myocyte size.

Protein expression of α1 and β1 subunits of the Na^+^-K^+^ pump and the loading control GADPH were measured from tissue homogenate. Protein was separated with SDS-PAGE, immunoblotted and then probed with corresponding antibodies.

### Patients

Patients had no valvular stenosis or regurgitation as the primary cause of HF and, as independently assessed by ≥2 cardiologists, HF was refractory to treatment short of i.v. inotropes or had relapsed after such treatment. Taking into account the adverse effects inotropes have on long-term outcomes ^14, 15^, most patients were not treated with these before mirabegron was prescribed.

We prescribed mirabegron off-label, permissible in Australia if in the opinion of the prescriber it is in the best interest of the patient and is supported by reasonable evidence ^16^. Approval by the institutional review board was not required. Consent for treatment and publication of de-identified case histories was obtained.

We gave mirabegron orally in a dose of 25 mg b.d. if the glomerular filtration rate (GFR) was at least 30 ml/min/1.73 m^2^. With a GFR <30-but ≥15 ml/min/1.73 m^2^ we reduced the dose, usually to 25 mg daily and did not use mirabegron at GFRs <15 ml/min/1.73 m^2^. Clinical data presented is observational, reflects the practice of the patients’ usual cardiologist and was not acquired according to pre-determined protocols.

### Statistics

Results are expressed as mean ± standard error (SE) unless indicated otherwise. Comparisons are made with analysis of variance (ANOVA) or paired t-test. A post-hoc Bonferroni test is used for multiple comparisons as appropriate. Comparisons for categorical variables are made with a Fisher’s exact test.

## Results

### Hemodynamic Variables and Clinical Indices of Heart Failure in Rabbits

Treatment with CL316,243 did not alter heart rate or blood pressure significantly in sham-operated rabbits or in rabbits with HF and there was no significant effect of HF or treatment with CL316,243 on serum creatinine (Table 1).

**TABLE 1.** Haemodynamic variables and serum creatinine in CL316,243 and ASP9531 treated sham- and congestive heart failure rabbits.

|  | Sham | Sham+CL | HF | HF+CL | HF | HF+ASP |
| --- | --- | --- | --- | --- | --- | --- |
| Rabbits | 10 | 8 | 13 | 8 | 6 | 6 |
| HR<br>(beats min<sup>-1</sup>) | 233 ± 2 | 230 ± 5 | 295 ± 15 | 268 ± 6 | 294 ± 22 | 316 ± 11 |
| MAP<br>(mmHg) | 79 ± 7 | 73 ± 4 | 70 ± 6 | 72 ± 9 | 72 ± 4 | 74 ± 7 |
| Creatinine<br>(μmol/L) | 76 ± 5 | 71 ± 4 | 90 ± 10 | 76 ± 9 | 94 ± 18 | 83 ± 15 |

Hemodynamic data and serum creatinine levels from sham, CL316,243 (CL) treated sham, heart failure (HF), CL and ASP9531 (ASP) treated HF groups. Result are presented as mean ± SEM.

Coronary ligation led to a severe HF phenotype with increased lung: body weight- (Figure 1A), heart: body weight- (Figure 1B) and liver: body weight ratios (Figure 1C). Most rabbits that had coronary ligation and were not treated with CL316,243 had ascites (Figure 1D). Sham-operated rabbits or sham-operated rabbits treated with CL316,243 did not have ascites (Figure 1D). Treatment with CL316,243 of rabbits that had coronary ligation reduced lung- (Figure 1A), heart- (Figure 1B) and liver weight and prevalence of ascites (Figure 1D).

**Figure 1.**
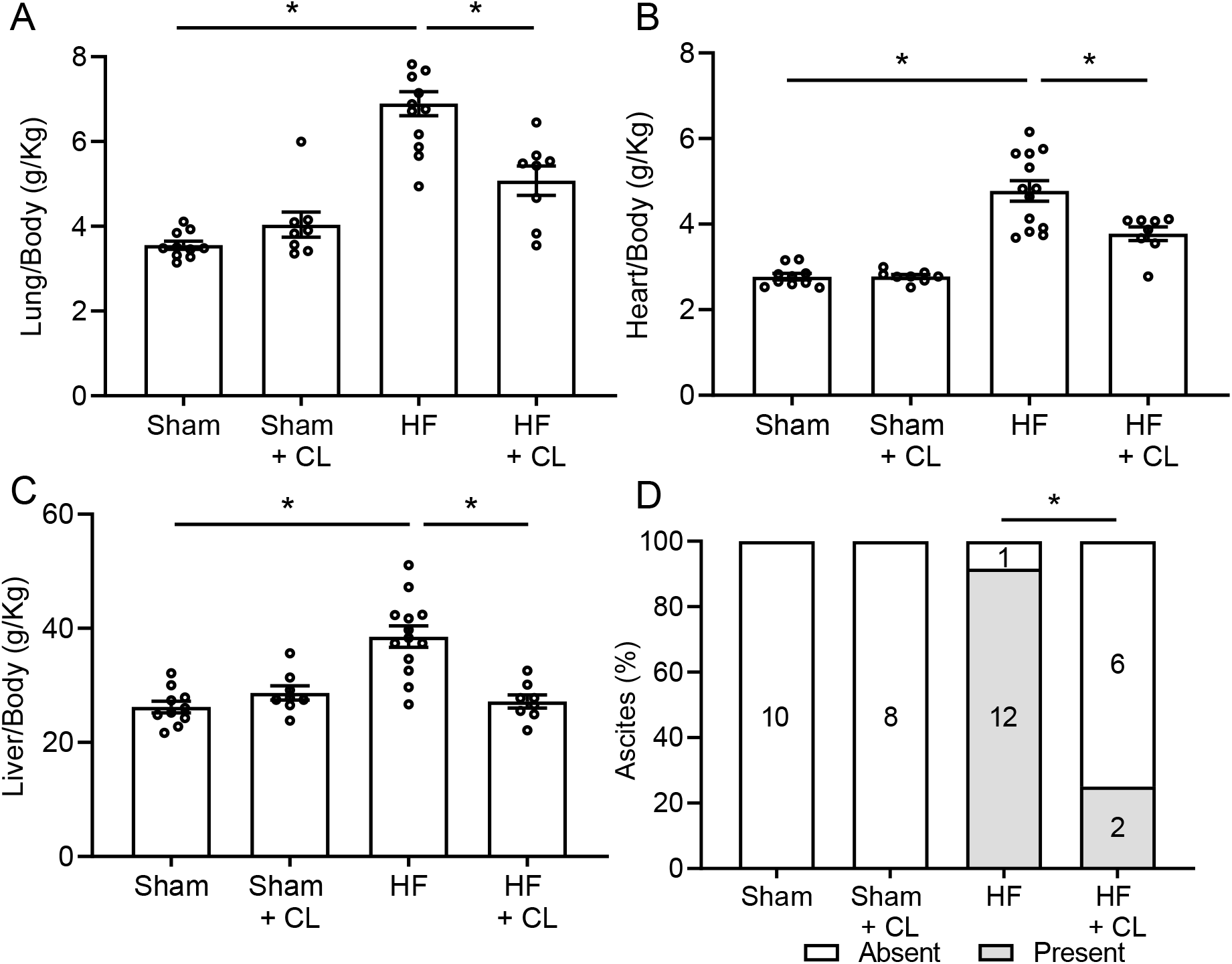
Indices of heart failure in coronary ligation model and effects of treatment with CL316,243. **A.** Lung: body weight ratio, **B.** Heart: body weight ratio, **C.** Liver: body weight ratio. **D.** Ascites in sham-operated rabbits and rabbits with heart failure (HF), with or without treatment with CL316,243 (CL). Numbers of rabbits are indicated in columns. Data in panels A, B and C are presented as mean ± SEM. P values were determined by ANOVA followed by Bonferroni multiple comparisons test. Data in panel D is compared using Fisher’s exact test. * indicates P < 0.05.

### Na^+^-K^+^ Pump Function in Heart Failure and Response to CL316,243 Treatment

Measurements of I_p_ in myocytes from a rabbit with HF and a rabbit with HF treated with CL316,243 are illustrated in Figure 2A. I_p_ was lower in myocytes from rabbits with HF than in myocytes from sham-operated rabbits. Treatment with CL316,243 eliminated the decrease in I_p_ with HF (Figure 2B). It also increased I_p_ of myocytes from sham-operated rabbits free of HF (Figure 2B) consistent with an increase in I_p_ after in vitro exposure to CL316,243 of myocytes isolated from normal rabbits that had not undergone surgery ^4^.

**Figure 2.**
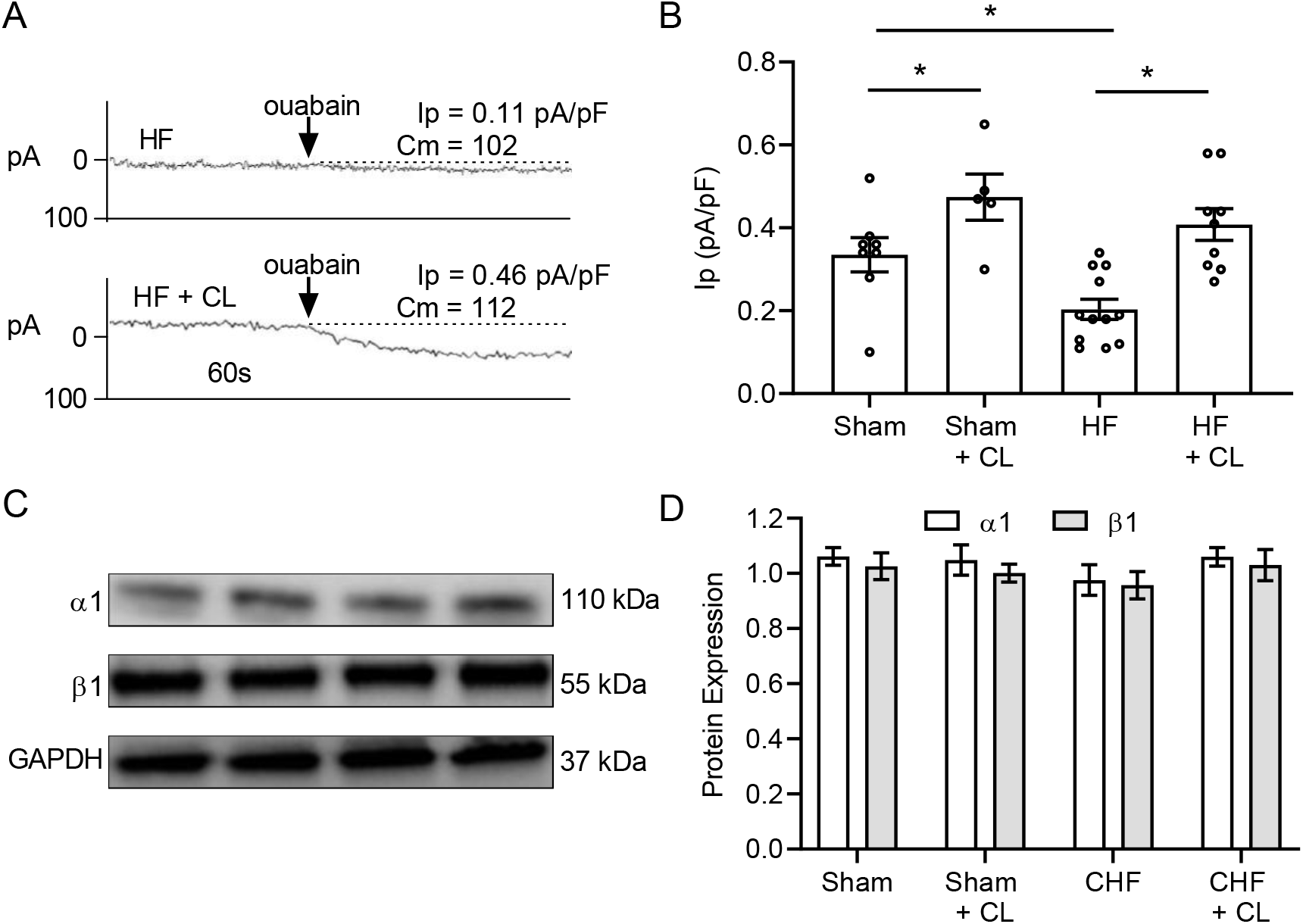
Effects of heart failure and treatment with CL316,243 on Na^+^-K^+^ pump function. **A**. Membrane currents of myocytes isolated from rabbits with CHF treated or not treated with CL. I_p_ was identified by the shift in the holding current induced by ouabain. C_m_ indicates membrane capacitance. **B**. Effects of CHF and treatment with CL on mean I_p_. Number of myocytes are indicated in the columns (isolated from ≥ 5 rabbits for each treatment group). * indicates P < 0.05. **C**. Representative blots of the effects of CHF and treatment with CL on expression of the α_1_ and β_1_ Na^+^-K^+^ pump subunit. GAPDH was used as the loading control. **D**. Effects of CHF and treatment with CL on expression of the α_1_ and β_1_ pump subunit. Expressions of the α_1_- and β_1_ pump subunits were measured for 6 rabbits in each treatment group. Data is presented as mean ± SEM. P values were determined by ANOVA followed by Bonferroni multiple comparisons test. * indicates P < 0.05.

The main Na^+^-K^+^ pump constituents are a large catalytic α subunit and a smaller β subunit with the α1 and β1 isoforms the most abundant in rabbit heart. There was no significant difference between expression of α1- (Figure 2C) or β1 subunits (Figure 2D) in myocardial homogenate from sham-operated rabbits or from rabbits with HF, with or without CL316,243 treatment, indicating that CL316,243 increased Na^+^-K^+^ pump turnover rather than abundance.

### Treatment with ASP9531

ASP9531 is highly specific for the human β3 AR. We ascertained it stimulates the Na^+^-K^+^ pump in rabbit cardiac myocytes in vitro as shown for other β3 AR agonists previously ^4^. Myocytes were isolated from normal rabbits as they were in the previous study ^4^. After establishing the whole-cell voltage clamp configuration, myocytes were exposed to 200 nmol/L ASP9531 for ~5 min before I_p_ was measured. Mean I_p_ was 0.52 ± 0.07 pA/pF for 6 myocytes exposed to ASP9531 and 0.30 ± 0.02 pA/pF for 8 control myocytes (P< 0.01).

We also measured I_p_ in myocytes from normal rabbits that had been given ASP9531 (30 mg/kg/24 hours) orally for 7 days. Mean I_p_ of myocytes isolated from 5 rabbits given ASP9531 was 0.47 ± 0.06 pA/pF and 0.30 ± 0.02 pA/pF in myocytes from 5 control rabbits (P <0.05). Oral treatment proved impractical (see Data Supplement) and for rabbits with HF we gave ASP9531 via osmotic minipumps.

Duration of treatment with ASP9531 was longer than that of treatment with CL316,243 (14 vs. 3 days) and there was therefore a longer time between coronary ligation of the rabbits and the time of their sacrifice (~28 vs. ~14 days). To limit the number of rabbits undergoing thoracotomy a sham-operated control group was not used again.

ASP9531 did not significantly alter heart rate, blood pressure or serum creatinine (Table 1). The lung: body weight ratio in rabbits with HF for 28 days (Figure 3A) was similar to that after 14 days (Figure 2A) suggesting that severity of HF was stable after day 14. Treatment with ASP9531 for 14 days reduced the lung: body- (Figure 3A) and heart: body weight ratios (Figure 3B) but there was no statistically significant effect on the liver: body weight ratio (Figure 3C - see comment in legend). Treatment eliminated ascites (Figure 3D).

**Figure 3.**
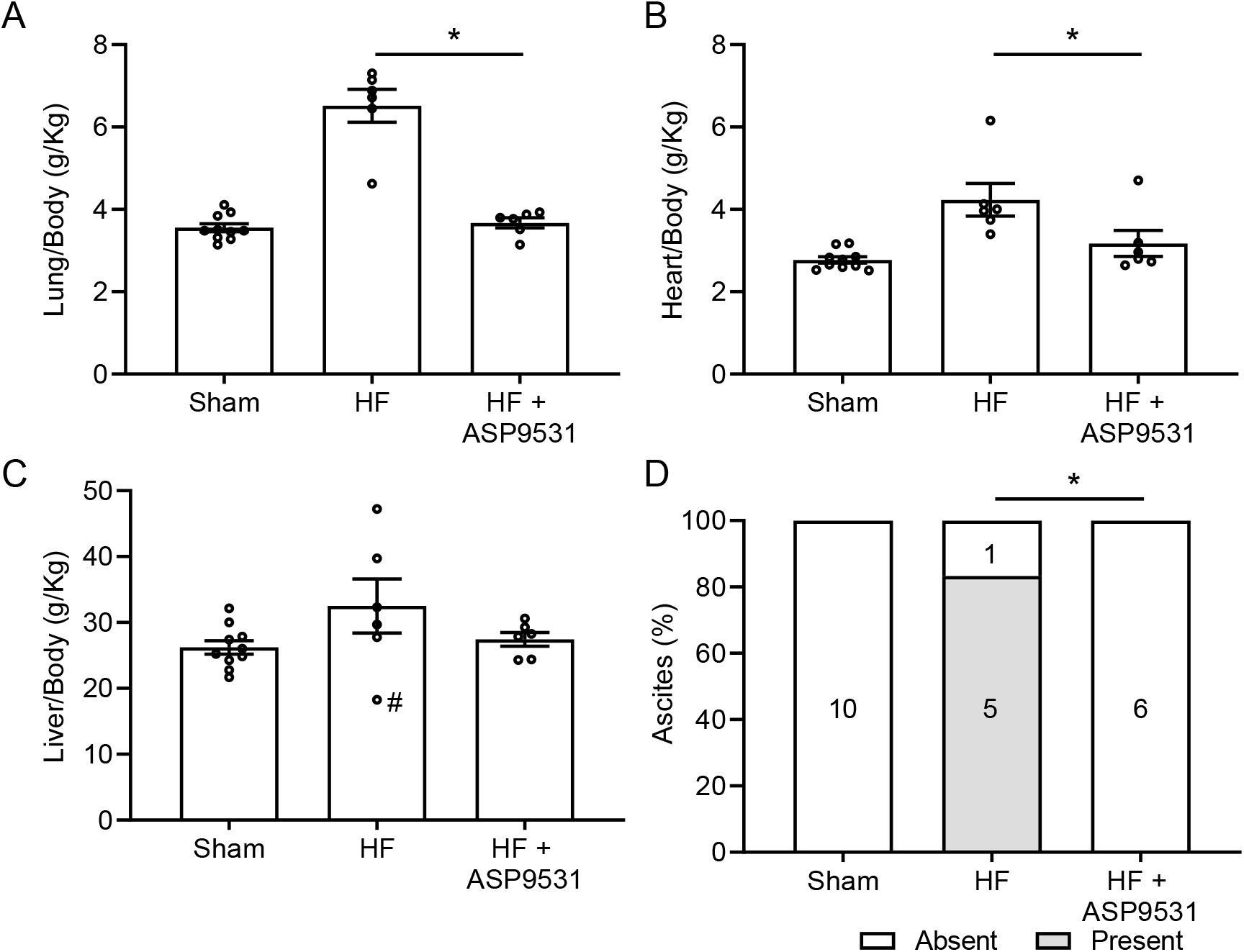
Indices of congestive heart failure with and without treatment with ASP9531. **A**. Effect of treatment with ASP9531 on lung: body weight ratio. Ratios for sham-operated rabbits are the same as those in Figures 1 A-C and are not used for statistical comparisons here. **B**. Heart: body weight ratios. **C**. Liver: body weight ratios. Data from a rabbit in the HF group with a congenitally small liver, reducing the mean ratio and increasing variance of ratios, is indicated by #. **D**. Prevalence of ascites at autopsy. Numbers of rabbits are indicated in columns. Data in panels A, B and C are presented as mean ± SEM. P values were determined by ANOVA followed by Bonferroni multiple comparisons test. Data in panel D is compared using Fisher’s exact test. * indicates P < 0.05.

### Mirabegron in severe treatment-resistant human HFrEF

Nine patients aged 74 ± 7 years (all clinical data is presented as mean ± standard deviation) had been admitted to hospital with worsening chronic heart failure (WCHF). Clinical characteristics of the patients, their treatments and outcomes are summarized in Table 2. Case histories are presented in the Data Supplement.

**Table 2.** Summary of Patient Characteristics and Outcomes

| Patient | 1 | 2 | 3 | 4 | 5 | 6 | 7 | 8 | 9 |
| --- | --- | --- | --- | --- | --- | --- | --- | --- | --- |
| Age (years) | 84 | 70 | 73 | 68 | 84 | 76 | 80 | 68 | 64 |
| Gender | Male | Male | Female | Male | Male | Male | Male | Male | Male |
| Diabetes | – | – | – | + | – | – | – | + | + |
| Ischaemic<br>Cardiomyopathy | + | – | – | + | + | + | – | + | + |
| Valvular disease | – | + | + | – | – | – | – | – | – |
| LVEF (%) | 35-40 | 28 | 20-25 | 25 | 25-30 | 25-30 | 20 | 30 | 20 |
| BNP (ng/L, < 159) | N/A | 1553 | 7005 | 882 | 991 | 1033 | N/A | 2642 | 2591 |
| NYHA Class | IV | III | IV | IV | III | IV | IV | III | IV |
| Hyponatremia | – | – | + | – | + | – | – | – | – |
| Previous inotropes | – | – | + | – | – | – | + | – | + |
| Bi-V pacing/ICD | – | +/+ | +/+ | +/+ | -/+ | – | +/+ | – | -/+ |
| Creatinine (μmol/L) | 135 | 89 | 85 | 70 | 131 | 95 | 221 | 240 | 310 |
| Follow-up (months) | 30* | 27* | 42 | 2* | 40 | 37 | 16† | 32 | 10* |
| Furosemide before‡¶ | 120 | 160§ | 80 | 480§ | 100§ | 160§ | 120 | 40 | 240§ |
| Furosemide after‡¶ | 40 | 100 | 40 | 240 | 40 | 80 | 80 | 60 | 160 |
| Spironolactone before‡¶ | 12.5 | 25 | 25 | 50 | 25 | 12.5 | 25 | – | 25 |
| Spironolactone after‡¶ | 12.5 | 25 | 25 | 50 | 25 | 12.5 | 25 | – | 25 |
| Thiazide diuretic | – | – | – | + | – | – | – | – | + |
| Beta blocker | – | + | + | + | – | + | + | + | + |
| ACE inhibitor before | + | – | – | + | – | – | + | – | – |
| ACE inhibitor after | – | – | – | + | – | – | – | – | – |

\* Deceased
† Deceased from progressive heart failure
‡ Cumulative daily dose in mg
§ Maximal oral equivalent of iv administration assuming 50% bioavailability with oral dosing

A cause precipitating cardiac decompensation of the patients had not been identified. They had been treated for HFrEF for ≥ 6 months. All had symptoms and clinical signs of pulmonary congestion or edema. Eight had leg and/or sacral edema and 5 had abdominal distension attributed to ascites. Three were in New York Heart Association (NYHA) Class III and 6 in NYHA Class IV and all had advanced, stage D HF ^7, 10^. BNP, available for 7, was 2385 ± 2167ng/L.

Four patients had bi-ventricular pacing and an implantable cardiac defibrillator (ICD); two had a defibrillator alone but not bi-ventricular pacing. The most recent LVEF measured by echocardiography was 26 ± 5%. The systolic blood pressure before treatment with mirabegron was 110 ± 13 mm Hg. This value includes the blood pressures of 2 patients while they had a second course of i.v. inotropic support when early relapse of severe de-compensation had occurred after a first course. Mirabegron was introduced during a bridging period before inotropes were weaned off in these patients. Their serum creatinine was raised to 221 and 310 μmol/L before they were given the second course of inotropes. The inotropes reduced these to levels that allowed use of mirabegron. The averaged serum creatinine for all 9 patients, including the 221 and 310 μmol/L levels, was 153 ± 84 μmol/L. One patient had inotropic support during an admission 4 weeks previously but was not given inotropes during the index admission.

HF had been refractory to escalating treatment that included i.v. furosemide unless HF was complicated by hyponatremia or renal function was deemed compromised by intravascular hypovolemia. Eight patients were also treated with spironolactone before mirabegron was introduced but only 1 at recommended target dose. Two patients were treated with a thiazide diuretic. ACE inhibitors had been discontinued in 7 patients due to low blood pressure and/or renal impairment. One had an ACE inhibitor replaced with sacubitril/losartan in hospital. This caused marked symptomatic hypotension needing immediate treatment with i.v. saline. Sacubitril/losartan was discontinued before mirabegron was introduced. β1 AR antagonists had been prescribed in 8 patients but for 7 at a dose ≤50% of recommended target.

There was no decrease in blood pressure with the addition of mirabegron in any patient, physical signs and symptoms improved and all were stabilized and discharged from hospital. This was achieved with a lower daily dose of furosemide at the time of discharge (median 80 mg) than at the start of treatment with mirabegron (median 120 mg, p < 0.05, paired t-test). Three patients aged 68, 73 and 76 years resumed full- or part time work.

Five patients died 17 ± 13 months after starting treatment with mirabegron. One was readmitted with HF 10 months after discharge and subsequently died from heart failure 16 months after treatment with mirabegron was started. Four died from other causes 2 – 30 months (median 18.5 months) after discharge. One of them died at 30 months from pneumonia that he declined treatment for because co-morbidities markedly limited his quality of life. Two have died from cancer. One died from a cardiac arrest 10 months after treatment with mirabegron was initiated. He had diabetic kidney disease that progressed to end-stage. In contrast to what had been the case during his index admission, renal function had not responded to i.v. inotrope treatment. He had required hemodialysis and mirabegron had been discontinued before he died.

Four patients remain alive 38 ± 4 months after treatment with mirabegron was introduced. While co-morbidities for some patients preclude accurate assessment of cardiac limitation, improvement ≥ 1 NYHA Class was maintained post-discharge for all except for the patient who died from heart failure 16 months after starting treatment with mirabegron and the patient who had progressive kidney disease. Outcomes after discharge from hospital are summarized in Figure 4.

**Figure 4.**
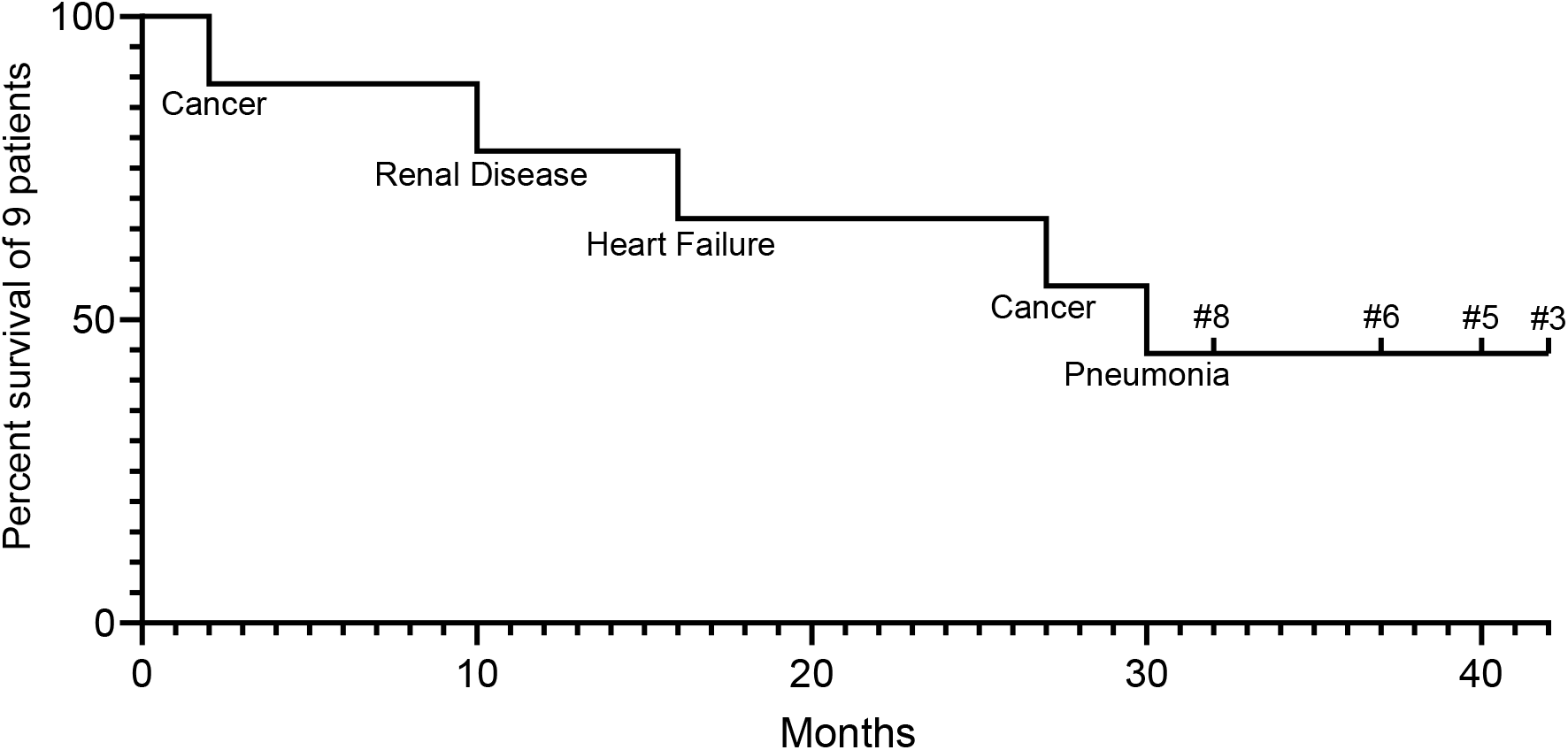
Survival of patients after mirabegron was added to their treatment. Each step indicates the death of a patient with the primary cause of death also shown. Numbers #8, #6, #5 and #3 of surviving patients refer to numbering of patients in Table 2.

## Discussion

While a modest increase of [Na^+^_i_] above baseline increases contractility of normal human myocardium in vitro, there is an inverse relationship between [Na^+^_i_] beyond already raised levels and contractility in myocardium from patients with severe HFrEF ^3^. This increases [Ca^2+^_i_] due to reduced Ca^2+^_i_ export via Na^+^-Ca^2+^ exchange, and dysregulation of Ca^2+^_i_-dependent excitation-contraction coupling causes contractile dysfunction ^17^. A high [Na^+^_i_] also causes mitochondrial dysfunction and energy deficiency ^18^.

Raised [Na^+^_i_] in the myocardium of patients with severe HFrEF is in part due to influx through the late Na^+^ current ^19^. Reduced Na^+^-K^+^ pump-mediated Na^+^_i_ efflux, here estimated by ~40% in rabbit myocytes with [Na^+^_i_] clamped at 10 mmol/L, also contributes. Direct measurement of [Na^+^_i_] and the relative contributions the late Na^+^ current and Na^+^-K^+^ pump inhibition make to an increase of [Na^+^_i_] in the beating heart is not experimentally possible. However, from in silico modelling ^20^, we estimate a decrease in Na^+^-K^+^ pump-mediated Na^+^_i_ efflux by 40% contributes much more than increased Na^+^ influx *via* the late Na^+^ current.

Raised [Na^+^_i_] could become self-sustaining because it stimulates mitochondrial synthesis of reactive oxygen species (ROS) ^21^ that in turn can activate the late Na^+^ current ^22^ and inhibit the Na^+^-K^+^ pump ^23^. Na^+^-K^+^ pump inhibition is countered by activation of the β3 AR that mediates antioxidant effects via the classical NO-dependent signalling pathway ^24^. This is experimentally supported by the reversal of pump inhibition caused by oxidative stress with in vitro exposure of cardiac myocytes to CL316,243 ^4, 23^. The reversal should mitigate Na^+^_i_-dependent Ca^2+^_i_ dysregulation and energy deficiency and hence contractile dysfunction.

Parallel reversal of Na^+^-K^+^ pump inhibition and organ congestion in rabbits treated with CL316,243 is in keeping with the hypothesis that cellular and molecular abnormalities in severe HFrEF are potential treatment targets ^1, 2^ and with an inhibited Na^+^-K^+^ pump as one such target. However, Na^+^-K^+^ pump inhibition might not have remained a useful target if CL316,243 had been added to treatment with drugs used in guideline-directed clinical practice because the most commonly used of these prevent Na^+^-K^+^ pump inhibition caused by angiotensin II receptor- ^23^ β1 AR- ^23^ and aldosterone receptor activation ^25^, effectively causing Na^+^-K^+^ pump stimulation, as reviewed ^5^.

The patients we report were given guideline-directed device therapy and drugs as tolerated before mirabegron was added. Not all recommended drugs were tolerated and of those prescribed most were not tolerated in target doses. With incomplete reversal of Na^+^-K^+^ pump inhibition it is likely myocardial [Na^+^_i_] was raised. This is directly supported by an abnormally high [Na^+^_i_] measured in the myocardium from patients with end-stage HFrEF. Although use of ACE inhibitors, β1 AR- and aldosterone receptor antagonist was not reported ^3^, it is reasonable to assume they were prescribed as tolerated. Patients most in need might be the least likely to tolerate evidence-based treatments that counter Na^+^_i_ accumulation ^5^, contributing to rapid decline and high mortality rates once HF reaches the advanced stage.

While sharing a putative raised [Na^+^i] as a treatment target with mainstay conventional drugs, β3 AR agonists-induced Na^+^-K^+^ pump stimulation actively reverses an increase in [Na^+^_i_] regardless of source ^9^ rather than incompletely countering inhibition of pump-mediated Na^+^_i_ export that the conventional drugs do. Consistent with this, treatment of patients with mirabegron was associated with improved physical signs and symptoms on a time scale compatible with the rapid resolution of Na^+^-K^+^ pump inhibition and congestive features in the rabbit model.

β3 AR agonist-induced Na^+^-K^+^ pump stimulation depends on activation of “soluble” guanylate cyclase (sGC) that is part of the classical NO-dependent signaling pathway ^4^. A direct activator of sGC stimulates the pump in cardiac myocytes ^13^ and activators of sGC have been trialed in HFrEF. Intravenous treatment of acute cardiac decompensation with cinaciguat improved hemodynamic indices but frequently caused hypotension and a trend towards increased re-admission rate ^26^. Oral treatment with vericiguat for 12 weeks of patients who were clinically stable after an episode of WCHF did not achieve the primary end point of a decrease in N-Terminal proBNP but was well tolerated. An exploratory secondary analysis suggested it might be beneficial ^27^ and an event-driven phase III trial is now in progress ^28^.

While β3 AR agonists and sGC activators share downstream signaling pathways, they are not expected to have identical effects in HF. Expression of sGC is virtually ubiquitous and it has a multitude of downstream effects ^29^. However, the β3 AR is differently expressed and coupled to cellular signaling domains in different cells and organs ^24^. In addition, myocardial expression of the β3 AR is up-regulated in end-stage human HFrEF ^30^ and, in contrast to the other β ARs, does not desensitize to agonist activation ^31^. These characteristics allow myocardial targeting in HF with greater selectivity than with activators of sGC and with fewer off-target effects.

Treatment of our patients with mirabegron off-label in doses recommended for the approved indication was well tolerated but might not be optimal for treatment of HF and up to 6-fold higher doses have been used in studies on overactive bladder ^32, 33^. There were no effects expected to be adverse in HF and consistent with this, none were apparent when patients with mostly NYHA Class II HFrEF were given the 6-fold higher dose ^9^. Mirabegron appears safe across the spectrum of severity of HFrEF.

A study is in progress on patients with HFrEF and NYHA Class III and IV. Effects of mirabegron on acute hemodynamics and remodelling after 3 months is being examined ^34^. Relevant to HF with preserved ejection fraction (HFpEF), that also can progress to advanced HF ^10^, a study on the effect of mirabegron on cardiac hypertrophy is in progress. Cardiac hypertrophy has a raised myocyte [Na^+^_i_] ^35^ as a treatment target in common with HFrEF and the study is examining if mirabegron improves LV hypertrophy and diastolic function in patients at high risk of developing HFpEF ^36^.

Here, there was a close temporal relationship between a favourable change in the clinical trajectory and treatment with mirabegron. Our studies on rabbits showed mechanistic plausibility for treatment efficacy and there was coherence with evidence-based therapies for HFrEF by the mechanistically shared stimulation of myocardial Na^+^_i_ export ^5^. These features suggest treatment of the patients with mirabegron might be efficacious ^37^ but do not prove it.

To our knowledge, only the REMATCH trial, that compared medical treatment with MCS ^8^, provides insight into long term outcomes of patients with advanced HF. The mainstay of evidence-based medical treatments for the patients recruited to the trial ~20 years ago has not changed substantially since then and the cardiovascular outcome of the patients we describe here compares favourably with the 75% 1 year- and nearly 100% 2 year mortality of 61 medically managed patients in the trial who mostly died from HF. Nevertheless, efficacy of mirabegron of course cannot be implicated.

The prevalence of advanced HF among all patients with HF is estimated at 1-10% and expected to increase ^10^. Cardiac transplantation or MCS as destination therapy is recommend in the management of otherwise intractable cases. However, many patients are not eligible for these treatments and few options beyond supportive care are available to them. Continuous infusion of inotropes at home may provide end-of-life symptomatic relief but it probably also reduces survival ^7, 10^.

Treatment with mirabegron as an alternative to usual care of those not eligible for transplantation or MCS is attractive. Mechanistically a β3 AR agonist addresses the need to target an identified myocardial abnormality in HF ^1, 2^ as shown here. Clinically an agonist addresses the need to integrate acute-with long-term management ^6, 9^, it can be used with GFR reduced to 15 ml/min/1.73 m^2^ and, as found here and previously ^9^, it does not cause the decrease blood pressure that often limits use of guideline-directed treatments. We submit a randomized controlled trial should be considered.

## Acknowledgements

We thank Astellas Pharma, Tokyo, for the gift of ASP9531.

## References

1. Gheorghiade M, Larson CJ, Shah SJ, Greene SJ, Cleland JG, Colucci WS, Dunnmon P, Epstein SE, Kim RJ, Parsey RV, Stockbridge N, Carr J, Dinh W, Krahn T, Kramer F, Wahlander K, Deckelbaum LI, Crandall D, Okada S, Senni M, Sikora S, Sabbah HN, Butler J. Developing New Treatments for Heart Failure: Focus on the Heart. Circ Heart Fail. 2016;9

2. Bayeva M, Sawicki KT, Butler J, Gheorghiade M, Ardehali H. Molecular and cellular basis of viable dysfunctional myocardium. Circ Heart Fail. 2014;7:680–691

3. Pieske B, Maier LS, Piacentino V, Weisser J, Hasenfuss G, Houser S. Rate dependence of [Na^+^]_i_ and contractility in nonfailing and failing human myocardium. Circulation. 2002;106:447–453

4. Bundgaard H, Liu CC, Garcia A, Hamilton EJ, Huang Y, Chia KK, Hunyor SN, Figtree GA, Rasmussen HH. β3 adrenergic stimulation of the cardiac Na^+^-K^+^ pump by reversal of an inhibitory oxidative modification. Circulation. 2010;122:2699–2708

5. Liu CC, Fry NA, Hamilton EJ, Chia KK, Garcia A, Karimi Galougahi K, Figtree GA, Clarke RJ, Bundgaard H, Rasmussen HH. Redox-dependent regulation of the Na^+^-K^+^ pump: new twists to an old target for treatment of heart failure. J Mol Cell Cardiol. 2013;61:94–101

6. Cotter G, Cohen-Solal A, Davison BA, Mebazaa A. RELAX-AHF, BLAST-AHF, TRUE-AHF, and other important truths in acute heart failure research. Eur J Heart Fail. 2017;19:1355–1357

7. Fang JC, Ewald GA, Allen LA, Butler J, Westlake Canary CA, Colvin-Adams M, Dickinson MG, Levy P, Stough WG, Sweitzer NK, Teerlink JR, Whellan DJ, Albert NM, Krishnamani R, Rich MW, Walsh MN, Bonnell MR, Carson PE, Chan MC, Dries DL, Hernandez AF, Hershberger RE, Katz SD, Moore S, Rodgers JE, Rogers JG, Vest AR, Givertz MM, Heart Failure Society of America Guidelines C. Advanced (stage D) heart failure: a statement from the Heart Failure Society of America Guidelines Committee. J Card Fail. 2015;21:519–534

8. Rose EA, Gelijns AC, Moskowitz AJ, Heitjan DF, Stevenson LW, Dembitsky W, Long JW, Ascheim DD, Tierney AR, Levitan RG, Watson JT, Meier P, Ronan NS, Shapiro PA, Lazar RM, Miller LW, Gupta L, Frazier OH, Desvigne-Nickens P, Oz MC, Poirier VL, Randomized Evaluation of Mechanical Assistance for the Treatment of Congestive Heart Failure Study G. Long-term use of a left ventricular assist device for end-stage heart failure. N Engl J Med. 2001;345:1435–1443

9. Bundgaard H, Axelsson A, Hartvig Thomsen J, Sorgaard M, Kofoed KF, Hasselbalch R, Fry NA, Valeur N, Boesgaard S, Gustafsson F, Kober L, Iversen K, Rasmussen HH. The first-in-man randomized trial of a beta3 adrenoceptor agonist in chronic heart failure: the BEAT-HF trial. Eur J Heart Fail. 2017;19:566–575

10. Crespo-Leiro MG, Metra M, Lund LH, Milicic D, Costanzo MR, Filippatos G, Gustafsson F, Tsui S, Barge-Caballero E, De Jonge N, Frigerio M, Hamdan R, Hasin T, Hulsmann M, Nalbantgil S, Potena L, Bauersachs J, Gkouziouta A, Ruhparwar A, Ristic AD, Straburzynska-Migaj E, McDonagh T, Seferovic P, Ruschitzka F. Advanced heart failure: a position statement of the Heart Failure Association of the European Society of Cardiology. Eur J Heart Fail. 2018;20:1505–1535

11. Audigane L, Kerfant BG, El Harchi A, Lorenzen-Schmidt I, Toumaniantz G, Cantereau A, Potreau D, Charpentier F, Noireaud J, Gauthier C. Rabbit, a relevant model for the study of cardiac beta 3-adrenoceptors. Exp Physiol. 2009;94:400–411

12. Tagami T, Kushimoto S, Yamamoto Y, Atsumi T, Tosa R, Matsuda K, Oyama R, Kawaguchi T, Masuno T, Hirama H, Yokota H. Validation of extravascular lung water measurement by single transpulmonary thermodilution: human autopsy study. Crit Care. 2010;14:R162

13. William M, Vien J, Hamilton E, Garcia A, Bundgaard H, Clarke RJ, Rasmussen HH. The nitric oxide donor sodium nitroprusside stimulates the Na^+^-K^+^ pump in isolated rabbit cardiac myocytes. J Physiol. 2005;565:815–825

14. Francis GS, Bartos JA, Adatya S. Inotropes. J Am Coll Cardiol. 2014;63:2069–2078

15. Mortara A. Inotropes and vasopressors in acute heart failure, when the devil dresses as an angel. Eur J Heart Fail. 2018;20:342–344

16. Day R. Off-label prescribing. Aust Prescr. 2013;36:5–7

17. Bers DM. Cardiac sarcoplasmic reticulum calcium leak: basis and roles in cardiac dysfunction. Annu Rev Physiol. 2014;76:107–127

18. Bay J, Kohlhaas M, Maack C. Intracellular Na(+) and cardiac metabolism. J Mol Cell Cardiol. 2013;61:20–27

19. Sossalla S, Wagner S, Rasenack EC, Ruff H, Weber SL, Schondube FA, Tirilomis T, Tenderich G, Hasenfuss G, Belardinelli L, Maier LS. Ranolazine improves diastolic dysfunction in isolated myocardium from failing human hearts--role of late sodium current and intracellular ion accumulation. J Mol Cell Cardiol. 2008;45:32–43

20. Trenor B, Cardona K, Gomez JF, Rajamani S, Ferrero JM, Jr., Belardinelli L, Saiz J. Simulation and mechanistic investigation of the arrhythmogenic role of the late sodium current in human heart failure. PLoS One. 2012;7:e32659

21. Kohlhaas M, Liu T, Knopp A, Zeller T, Ong MF, Bohm M, O’Rourke B, Maack C. Elevated cytosolic Na^+^ increases mitochondrial formation of reactive oxygen species in failing cardiac myocytes. Circulation. 2010;121:1606–1613

22. Song Y, Shryock JC, Wagner S, Maier LS, Belardinelli L. Blocking late sodium current reduces hydrogen peroxide-induced arrhythmogenic activity and contractile dysfunction. J Pharmacol Exp Ther. 2006;318:214–222

23. Chia KK, Liu CC, Hamilton EJ, Garcia A, Fry NA, Hannam W, Figtree GA, Rasmussen HH. Stimulation of the cardiac myocyte Na^+^-K^+^ pump due to reversal of its constitutive oxidative inhibition. Am J Physiol Cell Physiol. 2015;309:C239–250

24. Balligand JL. Cardiac Salvage by Tweaking with Beta3-adrenergic Receptors. Cardiovasc Res. 2016;111:128–133

25. Mihailidou AS, Bundgaard H, Mardini M, Hansen PS, Kjeldsen K, Rasmussen HH. Hyperaldosteronemia in rabbits inhibits the cardiac sarcolemmal Na^+^-K^+^ pump. Circ Res. 2000;86:37–42

26. Erdmann E, Semigran MJ, Nieminen MS, Gheorghiade M, Agrawal R, Mitrovic V, Mebazaa A. Cinaciguat, a soluble guanylate cyclase activator, unloads the heart but also causes hypotension in acute decompensated heart failure. Eur Heart J. 2013;34:57–67

27. Gheorghiade M, Greene SJ, Butler J, Filippatos G, Lam CS, Maggioni AP, Ponikowski P, Shah SJ, Solomon SD, Kraigher-Krainer E, Samano ET, Muller K, Roessig L, Pieske B, Investigators S-R, Coordinators. Effect of Vericiguat, a Soluble Guanylate Cyclase Stimulator, on Natriuretic Peptide Levels in Patients With Worsening Chronic Heart Failure and Reduced Ejection Fraction: The SOCRATES-REDUCED Randomized Trial. JAMA. 2015;314:2251–2262

28. Armstrong PW, Roessig L, Patel MJ, Anstrom KJ, Butler J, Voors AA, Lam CSP, Ponikowski P, Temple T, Pieske B, Ezekowitz J, Hernandez AF, Koglin J, O’Connor CM. A Multicenter, Randomized, Double-Blind, Placebo-Controlled Trial of the Efficacy and Safety of the Oral Soluble Guanylate Cyclase Stimulator: The VICTORIA Trial. JACC Heart Fail. 2018;6:96–104

29. Derbyshire ER, Marletta MA. Structure and regulation of soluble guanylate cyclase. Annu Rev Biochem. 2012;81:533–559

30. Moniotte S, Kobzik L, Feron O, Trochu JN, Gauthier C, Balligand JL. Upregulation of beta(3)-adrenoceptors and altered contractile response to inotropic amines in human failing myocardium. Circulation. 2001;103:1649–1655

31. Liggett SB, Freedman NJ, Schwinn DA, Lefkowitz RJ. Structural basis for receptor subtype-specific regulation revealed by a chimeric beta 3/beta 2-adrenergic receptor. Proc Natl Acad Sci U S A. 1993;90:3665–3669

32. Vij M, Drake MJ. Clinical use of the beta3 adrenoceptor agonist mirabegron in patients with overactive bladder syndrome. Ther Adv Urol. 2015;7:241–248

33. Chapple CR, Dvorak V, Radziszewski P, Van Kerrebroeck P, Wyndaele JJ, Bosman B, Boerrigter P, Drogendijk T, Ridder A, Van Der Putten-Slob I, Yamaguchi O, Dragon Investigator G. A phase II dose-ranging study of mirabegron in patients with overactive bladder. Int Urogynecol J. 2013;24:1447–1458

34. Bundgaard H. Beta 3 Agonist Treatment in Heart Failure-2 (BEAT-HF II). 2019

35. Aksentijevic D, O’Brien BA, Eykyn TR, Shattock MJ. Is there a causal link between intracellular Na elevation and metabolic remodelling in cardiac hypertrophy? Biochem Soc Trans. 2018;46:817–827

36. Pouleur AC, Anker S, Brito D, Brosteanu O, Hasenclever D, Casadei B, Edelmann F, Filippatos G, Gruson D, Ikonomidis I, Lhommel R, Mahmod M, Neubauer S, Persu A, Gerber BL, Piechnik S, Pieske B, Pieske-Kraigher E, Pinto F, Ponikowski P, Senni M, Trochu JN, Van Overstraeten N, Wachter R, Balligand JL. Rationale and design of a multicentre, randomized, placebo-controlled trial of mirabegron, a Beta3-adrenergic receptor agonist on left ventricular mass and diastolic function in patients with structural heart disease Beta3-left ventricular hypertrophy (Beta3-LVH). ESC Heart Fail. 2018;5:830–841

37. Glasziou P, Chalmers I, Rawlins M, McCulloch P. When are randomised trials unnecessary? Picking signal from noise. BMJ. 2007;334:349–351

