## Supplementary material for "Targeting cardiac myocyte Na^+^-K^+^ pump function with β3 adrenergic agonist in rabbits and patients with severe congestive heart failure": Data Supplement

### EXPANDED METHODS

#### Rabbit studies

*Ethics*

This project was approved by the Royal North Shore Hospital Animal Care and Ethics Committee that operates in accordance with the NSW Animal Research Act (1985), Animal Research Regulation (2010) and the Australian code for the care and use of animals for scientific purposes (8th Edition 2013). Despite all care taken, suffering after thoracotomy cannot be fully eliminated, and protocols were designed to minimise the total number of rabbits used. Obtaining measurements of $I_p$ fulfilling quality criteria for acceptance [1, 2] is technically difficult and data on $I_p$ is not always obtained from a sacrificed rabbit. To reduce the risk of that, 2 - 3 researchers using separate experimental set-ups worked simultaneously on batches of myocytes isolated from the same rabbit. A total of 74 rabbits were used.

*Induction of heart failure*

Male New Zealand white rabbits (*Orytolagus cuniculus*), aged 14 weeks and weighing 2.5 – 3.6 kg were used, sourced from Flinders University School of Medicine Animal Facility, Adelaide, Australia. Subcutaneous buprenorphine hydrochloride (0.02 mg/kg) was administered one hour before surgical procedures. Anaesthetic induction was achieved with i.v. bolus injection of 2 mg/kg propofol via a marginal ear vein. Following immediate loss of consciousness, rabbits were intubated blindly with 3.0 mm uncuffed endotracheal tube. Anaesthesia was maintained with 2% isoflurane and an oxygen flow rate of 2 L/min. Oxygen saturation was monitored with a $SpO_2$ sensor placed on the ear. Adequate depth of anaesthesia was ensured by the absence of pedal and palpebral reflexes.

For haemodynamic measurements, a catheter was inserted into the central ear artery and connected to a blood pressure transducer (MLT0670 Blood Pressure Transducer; AD Instruments, Milford, MA. USA). Arterial pressure signals and heart rate during surgery were recorded (PowerLab/8sp; AD Instruments, Milford, MA. USA).

A thoracotomy was performed via the left 4$^{th}$ intercostal space. The pericardium was incised and the circumflex coronary artery ligated in its proximal part. Occlusion of the artery was ascertained by the development of cyanosis in the area at risk and ST segment elevation on the electrocardiogram monitored with leads on both forelimbs and left hind limb. After electrical and haemodynamic stability for >30 min a chest drain was placed at the 2$^{nd}$ intercostal space, the thoracotomy was closed in layers and the wound site sealed with an aerosol dressing spray. Haemo-pneumothorax was aspirated and when a negative intrathoracic pressure was restored the drain was removed. Sham operations were identical except that the coronary artery was not ligated.

Prophylactic cephalexin monohydrate, 30 mg/kg, was given i.v. after the surgical procedures and isoflurane administration was gradually decreased. The endotracheal tube was removed when coughing was observed. Rabbits were then returned to animal housing for recovery and postoperative analgesia (0.02 mg/kg buprenorphine hydrochloride given by subcutaneous injection every 8 hours for 48 hours). Two were found dead in the first week following ligation of the coronary artery early in the study. No deaths before rabbits were sacrificed according to protocol occurred subsequently.

At the time of sacrifice, rabbits were anesthetised with a subcutaneous injection of 80 mg/kg ketamine (Troy Laboratories Pty Ltd, Australia) and 50 mg/kg of ilium xylazine (Troy Laboratories Pty Ltd, Australia). Heparin, 1000 IU, was given through the marginal ear vein to prevent thrombus formation in coronary arteries. Deep anaesthesia was confirmed using pain stimulus induced by clamping the metacarpal space on hind and front paws and ascertaining absence of changes in breathing rate and absence of a corneal reflex. A sub-xyphoid incision was made and a median sternotomy performed. The aorta was then transected and the heart excised.

*Administration of β3 adrenergic agonists*

CL316,243 was administered using micro-osmotic pumps (Model 1003D, ALZET®, DURECT Corporation, Cupertino, CA, USA) with a mean pumping rate of 1.02 µl per hour to achieve an infusion rate of 2 mg/kg/24 hours. CL316,243 was dissolved in double distilled water immediately before filling of the pump. With this rate of *in vivo* infusion, electrogenic $Na^+$-$K^+$ pump current ($I_p$) measured in isolated rabbit ventricular myocytes measured *ex vivo* is increased as reported previously [3].

To implant micro-osmotic pumps, we induced anaesthesia with 2 mg/kg propofol i.v. and then maintained anaesthesia with 2% isoflurane and an oxygen flow rate of 2 L/min given via a facial mask. Using aseptic technique, a pocket was made *via* an incision between the scapulae. After pumps were inserted the pocket was closed with sutures and rabbits were recovered from anaesthesia as described above.

As ASP9531 is not water soluble we initially administered it in the oral dose of 30 mg/kg/24 hours that was recommended by the manufacturer for rabbits. We administered it by mixing it with honey which, when smeared on the fur, was licked off by the rabbits. We measured $I_p$ in myocytes from 5 rabbits given ASP9531 in this manner and from 5 control rabbits not given ASP9531. However, it was difficult to ascertain the dose received. In addition, rabbits appeared distressed when honey was repeatedly placed on their fur. Osmotic pumps (Model 2ML2, ALZET®, DURECT Corporation, Cupertino, CA, USA) with a mean pumping rate of 5 µl per hour were then used. We dissolved ASP9531 in 50% DMSO and double distilled water immediately before filling pumps according to the manufacturers' instructions to achieve a rate of administration of 250 µg/kg/24 hours.

*Echocardiographic assessment of left ventricular function*

Echocardiography (Toshiba Xario Ultrasound with 6.5 with PST-65AT 6.5 6.5 MHz paediatric transducer) was performed in anaesthetised rabbits before coronary artery ligation or sham operations, 7 days after surgery and again after the treatment period, just before sacrifice. Two-dimensional and M-mode measurements were made at short axis basal, mitral and apex levels, and with a long axis view. The LV ejection fraction (LVEF) was derived from end diastolic- and end systolic volumes calculated according to a modified Simpson's rule [4] to identify rabbits with heart failure (HF). Five rabbits had a LVEF > 25% after coronary ligations and were not allocated to any control or treatment group. In view of the segmental nature of LV dysfunction after coronary ligation, we considered variance of echocardiographic measurements too high for them to be suitable as an outcome variable. In addition, the LVEF does not necessarily reflect the congestive features that are targeted for treatment in clinical practice. The lung: body weight ratio measured at autopsy was the primary outcome variable. Liver: body weight- and heart: body weight ratios were also recorded.

Eighteen rabbits that had a sham operation and 33 rabbits that had had coronary ligation and had an ejection fraction of 25% or less after 7 days were assigned to treatment or no treatment with CL316,243 or ASP9531. Two additional animals that died are not reported on.

*Measurement of $Na^+$-$K^+$ pump current in voltage clamped ventricular myocytes*

Myocytes isolated from non-infarcted parts of the myocardium were voltage clamped using the whole-cell voltage clamp technique for measurement of electrogenic $Na^+$-$K^+$ pump current ($I_p$, arising from the 3$Na^+$: 2$K^+$ exchange ratio). Wide-tipped (4-5 μm) patch pipettes were filed with solution containing (in mM): 70 $K^+$-glutamate, 1 $KH_2PO_4$, 5 HEPES, 5 EGTA, 2 MgATP, 10 $Na^+$-glutamate, 80 tetramethylammonium chloride (TMA-Cl). The solution was titrated to a pH of 7.22 at 22°C using 1 M KOH. Myocytes were initially superfused with modified Tyrode's solution containing (in mM) 140 NaCl, 5.6 KCl, 2.16 $CaCl_2$, 0.44 $NaH_2PO_4$, 10 glucose, 1.0 $MgCl_2$ and 10 HEPES, titrated to a pH of 7.4 at 35°C with NaOH. After the whole-cell configuration was established, we switched to a superfusate that was nominally $Ca^{2+}$ free and contained 0.2 mM $CdCl_2$ and 2 mM $BaCl_2$. $Cd^{2+}$ was included to block $Ca^{2+}$ channel conductance and inhibit $Na^+$-$Ca^{2+}$ exchange. $Ba^{2+}$ was included to block $K^+$ channels.

Myocytes were voltage-clamped at -40 mV to inactivate voltage-sensitive $Na^+$ channels and L-type $Ca^{2+}$ channels. $I_p$ was identified at 37 °C as the shift in stable holding current induced by 100 μM ouabain. Criteria for acceptance of accurate measurements of $I_p$ were published previously [1, 2].

To ascertain that ASP9531 directly stimulates the $Na^+$-$K^+$ pump *in vitro*, we dissolved it in 50% DMSO which was then diluted to a final concentration of 0.13% in the nominally $Ca^{2+}$-free superfusate. DMSO in a concentration of 0.4% has no effect on $Na^+$-$K^+$-pump current in measured in voltage clamped cardiac myocytes [5]. Cardiac myocytes from normal rabbits that had not had a thoracotomy were used. To limit the number of animals used, myocytes from the same rabbit were exposed or not exposed to ASP9531. A total of 6 rabbits were used.

*Chemicals and Reagents*
TMA.Cl was purum grade and obtained from Fluke Chemicals (Switzerland). Chemicals used in Tyrode's solutions were analytical grade and obtained from BDH (Australia) and Sigma Chemical Co (St Louis, MO). Ouabain and CL316,243 was obtained from Sigma Chemical Co (St Louis, MO, USA). ASP9531 was kindly donated by Astellas Pharma Inc. (Tokyo, Japan). Antibodies to α1- and β1 subunits of $Na^+$/$K^+$-ATPase were obtained from Millipore, Billerica, Maryland, USA and an antibody to GADPH was obtained from Sigma Chemical Co. St Louis, Missouri, USA.

*Estimation of number of rabbits needed for experiments*
$I_p$ measured with the whole-cell voltage clamp technique reflects $Na^+$-$K^+$ pump activity in an intact cell. The technique also controls membrane voltage, important because $Na^+$-$K^+$ pump activity is voltage-dependent [5]. However, without careful attention to technical detail, measurements of $I_p$ are inaccurate and/or misleading. It is critically important that control of the concentrations of intracellular $Na^+$-$K^+$ pump ligands are achieved and this requires use of wide-tipped patch pipettes and low series resistance once the whole-cell configuration is established [5, 6,]. The technical difficulty with this complicates study design when it is important to achieve objectives with the lowest number of rabbits undergoing a thoracotomy. Establishing the whole-cell configuration and obtaining measurements of $I_p$ that satisfy pre-determined quality criteria [1] is not always achieved in myocytes from a rabbit and this determined study design and number of rabbits used.

Using a coefficient of variation of 15% that is robust across many experimental models from our laboratory, 6 valid recordings were needed to detect a 25% decrease with 80% power and a two-tailed $P<0.05$ in $I_p$ in myocytes from rabbits with HF, and similarly 6 recordings were needed to detect a 25% increase with treatment of such rabbits. We wished to obtain these recordings from ≥5 rabbits in each treatment arm. Once at ≥6 recordings from ≥5 rabbits had been obtained no more rabbits were entered into a treatment arm, but data was obtained and used from rabbits already assigned treatment but not yet sacrificed. We therefore report >6 recordings for each of the 3 most important groups of rabbits.

*Observer bias*

To eliminate observer bias, ouabain-induced shifts in holding currents were assessed against criteria for acceptance before the membrane capacitance was derived and used to normalise the $I_p$ we report. The capacitance is highly variable between cells, cannot be anticipated and determining it from recordings is not susceptible to observer interpretation. Similarly, no observer interpretation is involved in weighing lungs, heart and liver or determining presence or absence of ascites.

### SUPPORTING INFORMATION

#### Treatment of patients with β3 adrenergic agonist

***Patient 1***

A 83 year old male with ischaemic cardiomyopathy and previous coronary by-pass surgery was admitted to hospital with dyspnea on exertion, orthopnea and ankle swelling. LVEF was estimated at 35-40% 5 years previously. He had atrial fibrillation with a rate of ~100 beats/min. The jugular venous pressure was elevated to the angle of the jaw and he had pitting ankle edema. A chest X-ray showed pulmonary congestion and a left pleural effusion. Treatment for HF consisted of furosemide 40 mg mane with oral $K^+$ supplementation. He had not tolerated a β-blocker due to asthma and an ACE inhibitor had been stopped due to low blood pressure. Treatment was intensified to i.v. furosemide 80 mg mane and 40 mg midday. Oral $K^+$ supplementation was replaced with spironolactone 12.5 mg mane.

The escalated treatment had no effect on signs or symptoms over the following 5 days. The patient was then prescribed mirabegron, 25 mg mane. The heart rate decreased from ~100 beats/min to ~85 beats/min within a day and symptoms and signs of HF improved but did not fully resolve. He has discharged 11 days after mirabegron was first prescribed on treatment that included furosemide 40 mg mane, spironolactone 12.5 mg mane and mirabegron 25 mg mane.

The patient was readmitted 6 months later. He had stopped taking mirabegron and had gained a large amount of weight. He had exertional dyspnea, orthopnea and paroxysmal nocturnal dyspnea (had to sleep sitting in a chair). The jugular venous pressure was raised to the angle of the jaw, peripheral edema extended proximal to knees, there was broken skin with extravasation of tissue fluid at the ankles and he had a large shin ulcer. He had basal pulmonary crepitations. Marked abdominal swelling was attributed to ascites. He had atrial fibrillation with a rate of ~105 beats/min and a blood pressure of 100/60 mm Hg. His serum creatinine was 135 µmol/L.

He was initially treated with an increase in furosemide dose from 40 mg mane to 80 mg mane, 40 mg midday given orally and the dose of spironolactone was increased from 12.5 to 25 mg mane. Mirabegron was re-introduced, now in a dose of 25 mg b.d. The heart rate decreased by ~15 beats/min and the systolic blood pressure increased to 120/70 mm Hg within 2 days of re-introducing mirabegron. Symptoms and signs of HF resolved within 5 days and the weight decreased by 7.5 kg. The 40 mg midday dose of furosemide was omitted before discharge. Soon after the morning dose was decreased to 40 mg and the dose of spironolactone decreased to 12.5 mg.

With the higher mirabegron dose the patient had no recurrence of symptoms or clinical signs of HF. Mobility was limited by spinal canal stenosis and residual deficits after a cerebrovascular accident that had occurred before he was treated with mirabegron. He subsequently had admissions to hospital after falls or pneumonia but never for HF. He was treated with 25 mg b.d. mirabegron for 30 months until he died from pneumonia that he elected not to be treated for in view of debilitating co-morbidities.

***Patient 2***

A 70 year old male with aortic valve replacement 22 years previously had developed HF 18 years later. His LVEF then was 20-25%. He had previously had a bi-ventricular pacing device/implanted cardiac defibrillator (Bi-V/ICD). Treatment included bisoprolol 10 mg mane, furosemide 40 mg mane, spironolactone 12.5 mg mane. Treatment with an ACE inhibitor had been stopped due to low blood pressure.

The condition had been stable for 4 years after HF was first diagnosed when doses of furosemide and spironolactone had to be increased to 80 mg and 25 mg mane. Despite the escalation of treatment he had to be admitted to hospital with worsening dyspnea on minimal exertion and orthopnea. The jugular venous pressure was raised to the angle of the jaw, he had bilateral edema to the mid-shin and basal pulmonary crepitations. The BNP was 1553 ng/L. The blood pressure was 110/70 mm Hg. His serum creatinine was 89 µmol/L.

Treatment with furosemide was changed to 40 mg b.d. given i.v., with no improvement after 7 days. Mirabegron 25 mg b.d. was then added. The furosemide treatment was changed to 80 mg mane and 40 mg midday, given orally, a lower dose-equivalent than the 40 mg b.d. that had been given i.v. The patient reported he "felt better than he had for weeks" after the second dose of mirabegron and after 4 days ankle edema was eliminated, the jugular venous pressure was only raised by 2-3 cm and pleural effusions on the chest x-ray were almost eliminated (Figure 1). He was discharged 5 days after treatment with mirabegron was started. At follow-up 4 weeks later clinical improvement was maintained and radiological evidence of pleural effusions had been eliminated. Treatment with mirabegron remained well tolerated.

Improvement of symptoms and signs were sustained for 24 months after mirabegron was started until the patient developed initially unexplained tachycardia at a rate of 120 beats/min. This caused rapidly progressive HF progressing to cardiogenic shock that was only reversed when an ectopic atrial tachycardia tracked by his pacemaker was identified and abolished with amiodarone. He subsequently died from widely disseminated lung cancer 27 months after treatment with mirabegron had been commenced.

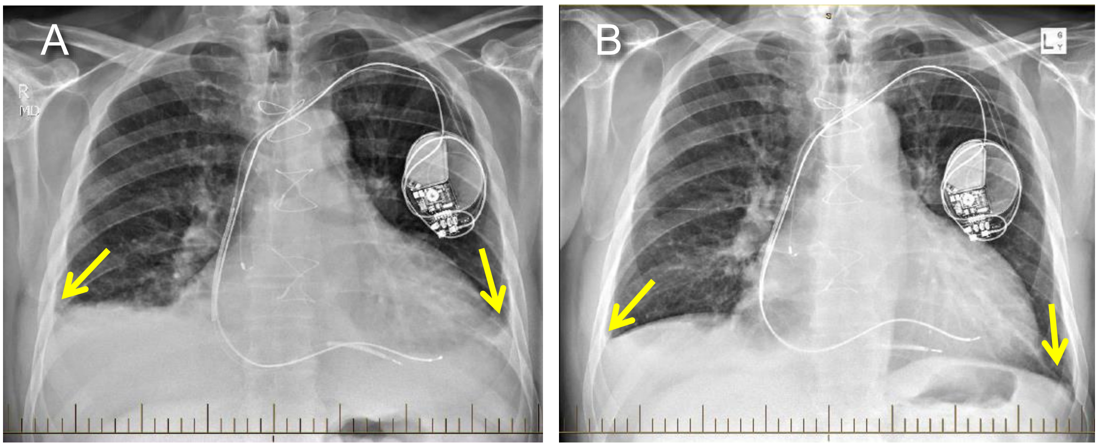


Figure 1. Chest X-rays of *Patient 2* taken 2 days before he commenced treatment with mirabegron (A) and 4 days after days after treatment was commenced. Large pleural effusions (arrows in panel A) were largely eliminated after treatment (panel B).

***Patient 3***

A 73 years old female, first diagnosed with dilated cardiomyopathy 10 years previously, had stable NYHA Class I-II HF on treatment with an ACE inhibitor and a β blocker when she presented with severe decompensated bi-ventricular failure complicated by hyponatraemia.

She had aortic regurgitation of mild-moderate severity on echocardiography and a LVEF of 20%. She also had severe chronic obstructive pulmonary disease. There was a good response to treatment with i.v. dobutamine and an iron infusion. She had a Bi-V/ICD implanted and was discharged from hospital with NYHA Class II symptoms.

Two weeks after discharge the patient developed worsening dyspnea and a month after discharge she was re-admitted with NYHA Class IV symptoms. She was being treated with furosemide 40 mg b.d., spironolactone 12.5 mg b.d. and bisoprolol 10 mg mane at the time of admission. The blood pressure was 110/55 mm Hg. There had been a large increase in serum BNP to 7005 ng/L since the previous discharge. Serum creatinine was 85 µmol/L. The previous hyponatraemia made treatment with diuretics precarious and treatment with mirabegron 25 mg b.d. was started. The dose of furosemide was reduced to 20 mg b.d.

The day after starting mirabegron the patient had a gastrointestinal haemorrhage. There had been no decrease in haemoglobin levels monitored daily before the haemorrhage that might have accounted for cardiac de-compensation. Endoscopy showed a gastric ulcer, attributed to use of aspirin. Blood transfusion restored haemoglobin to the level before the bleed. The gastric ulcer was managed conservatively and treatment with mirabegron continued.

In view of chronic obstructive pulmonary disease, bisoprolol was replaced with ivabradine with no change in heart rate. Symptoms improved to NYHA class II and serum BNP decreased to 588 ng/L before she was discharged. She returned to part-time work and has had no readmission for HF. Early follow up echocardiography showed a decrease in left ventricular dimensions and echocardiography 15 months after mirabegron was commenced showed a LVEF of 50%. She has maintained NYHA functional status class II for 42 months on continued treatment with mirabegron.

***Patient 4***

A 68 year old male had severe exacerbation HF which had become resistant to oral diuretic therapy. He had an ischaemic cardiomyopathy with a LVEF previously measured at 25%. A complex background included hypertension, anteroseptal myocardial infarction, coronary stenting, type 2 diabetes, pulmonary fibrosis secondary to chemotherapy for pancreatic cancer, ventricular tachycardia and an ICD with demand pacing. His medications were carvedilol 25 mg b.d., furosemide 120 mg b.d., ramipril 5 mg mane, ivabradine 7.5 mg b.d., spironolactone 25 mg b.d., hydrochlorothiazide 12.5 mg on alternate days, clopidogrel 75 mg daily and insulin.

He had lower limb edema extending to thighs and sacrum. His weight was 104 kg. The jugular venous pressure was elevated to the angle of the jaw. The blood pressure was 100/64 mm Hg. Serum BNP was 882 ng/L and serum creatinine was 70 µmol/L. The chest X-ray is shown in Fig. 2.

The patient was treated with i.v. furosemide 120 mg b.d. Ramipril was stopped and sacubitril/valsartan 24 mg/26 mg b.d. commenced. This caused hypotension with the blood pressure falling to 80/46 mmHg that was associated with symptoms of cerebral hypoperfusion and required emergency treatment with i.v. saline. Sacubitril/valsartan was discontinued.

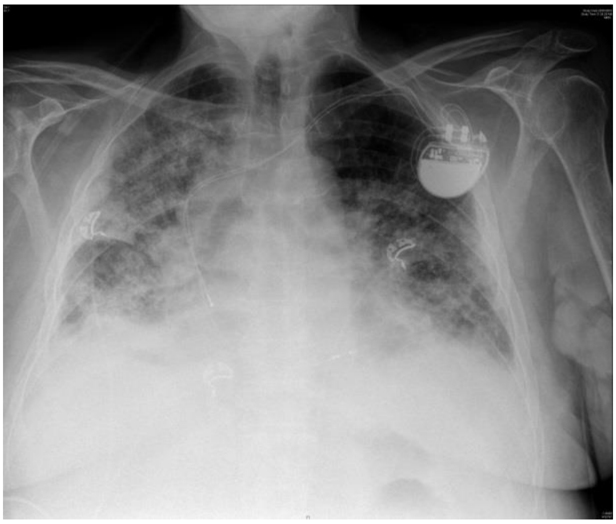

Figure 2. Chest X-ray for Patient 4 before treatment with mirabegron. There is extensive pulmonary fibrosis, thickening of the right fissure and small bilateral pleural effusions.

After a week of i.v. diuretic therapy, 2 weeks after admission to hospital, there was no change in weight or extent of edema. Mirabegron 25 mg b.d. was commenced as an alternative to i.v. dobutamine. Within 24 hours of the 2nd dose of mirabegron he reported feeling better and a diuresis began. Intravenous furosemide was stopped and oral therapy re-commenced. By day 7 after treatment with mirabegron had been started the weight had decreased by 6 kg and he was discharged home to the care of a community-based palliative care team.

On review by his cardiologist 2 weeks after discharge, the weight was 90 kg, 14 kg lower than before treatment with mirabegron. He had no edema and the jugular venous pressure was not elevated. The serum creatinine had increased to 132 µmol/L and his diuretic therapy was reduced. He continued to need oxygen therapy because of his pulmonary fibrosis but was able to stay at home until he died after a steady decline in his general condition 7 weeks after treatment with mirabegron was commenced. No shock had been delivered by the ICD at the time of death. An autopsy was not performed. Due to drowsiness, medication could only be given inconsistently the last week before he died and some lower leg edema had re-developed.

***Patient 5***

A 84 year old male had a myocardial infarction and coronary by-pass surgery 17 years previously. He had ischaemic cardiomyopathy and a Bi-V/ICD had been implanted 5 years previously at a time when his LVEF measured by echocardiography was 25-30%. Management was complicated by renal impairment with a serum creatinine of 131 µmol/L limiting use of diuretics. Treatment with ACE inhibitor had been stopped for the same reason. Bi-V pacing had improved his condition but leg edema had never been fully eliminated and had been present for several years. At times it had been associated with shin ulcerations. He had mild cognitive impairment secondary to multi-infarct strokes.

The patient was admitted to hospital with dyspnea at rest and peripheral edema extending proximal to the knees. The blood pressure was 135/75 mm Hg, serum BNP 991 ng/L and LVEF 25%. Symptoms had worsened despite a recent increase in the dose of furosemide from 40 mg mane and 20 mg at midday to 60 mg mane and 40 mg at midday. Other medications included amiodarone 200 mg mane for treatment of atrial fibrillation, spironolactone 12.5 mg b.d. and nebivolol 1.25 mg mane. Hyponatremia (123 mmol/L) had developed and treatment with diuretics and amiodarone was stopped and fluid restriction, nominally at 1.2 l/day introduced. Compliance with the fluid restriction was uncertain. There was no clinically detectable response to this treatment and mirabegron 25 mg b.d. was added to treatment 5 days after admission to hospital.

The degree of leg edema decreased and the patient lost ~7 kg in weight from the time of admission till discharge. Serum $Na^+$ levels increased and treatment with furosemide 40 mg mane and spironolactone 12.5 mg b.d. re-introduced. He tolerated up-titration of nebivolol to 7.5 mg mane and was discharged 13 days after admission, 8 days after he was first prescribed mirabegron. The nominal fluid restriction was lifted at the time of discharge.

At outpatient review 2 weeks later he remained free of dyspnea but some ankle edema remained. His serum $Na^+$ had normalised to 140 mmol/L despite the re-introduction of diuretics. At review 6 weeks after discharge he remained free of dyspnea, he had no leg/ankle edema nor did he have other physical signs of HF. He has not been readmitted to hospital for HF for 40 months on maintained treatment with mirabegron, hyponatremia has not recurred and there has been no increase in serum creatinine.

***Patient 6***

A 76 year old male had an anterior myocardial infarction treated with emergency angioplasty and placement of a stent to the left anterior descending coronary artery. This occurred on the background of rheumatoid arthritis treated with methotrexate with good control of disease.

The patient developed left ventricular dysfunction with a LVEF of 35% after the infarct. He was treated with carvedilol 3.125 mg b.d. at the time of discharge. An ACE inhibitor was not prescribed due to low blood pressure (systolic pressures 90 – 95 mm Hg). He re-presented to hospital 1 month post-discharge with dyspnea at rest. A chest X-ray showed pulmonary congestion. The LVEF was now 25 – 30%. Furosemide 40 mg mane was added to treatment and since asymptomatic, non-sustained ventricular tachycardia was detected on routine monitoring on a background of the reduced LVEF an ICD was implanted. His symptoms improved and he was discharged on carvedilol 3.125 mg b.d., furosemide 40 mg mane and spironolactone 12.5 mg mane. Attempts at up-titrating carvedilol dose during follow-up were not successful due to persistently low blood pressures, ~90 – 95 mm Hg systolic. He also developed weight loss that started soon after the initial presentation.

On a third admission, the patient presented at midnight with florid pulmonary edema (Figure 3A) and raised jugular venous pressure. His serum creatinine was 95 µmol/L. Symptoms improved after treatment with i.v. furosemide, topical nitrates and continuous positive airway pressure *via* a face mask. In view of the cardiac de-compensation despite optimised standard treatment the preceding months he was prescribed mirabegron 25 mg b.d. after cardiology review in the morning. He received his first dose at 10 AM and was then transferred from the emergency department to a general cardiology ward.

By midnight, pulmonary edema re-developed with radiological features more marked than on initial presentation (Figure 3B). Emergency treatment as above was repeated. It was realised that he had not received the prescribed evening dose of 25 mg mirabegron (was not in stock on the ward). The omitted dose was then given. The patient's family had been called in because of his critical condition and told by the emergency response team that "if this doesn't work there is nothing more we can do".

The patient improved gradually the following days and i.v. furosemide 40 mg b.d. was changed to oral administration of the same dose. His chest X-ray 5 days after the second episode of severe pulmonary edema showed marked improvement (Figure 4). His LVEF was 35% at the time of discharge and compared with the previous study 4 months previously (LVEF 25 – 30%) "there was improvement in apicolateral wall contractility".

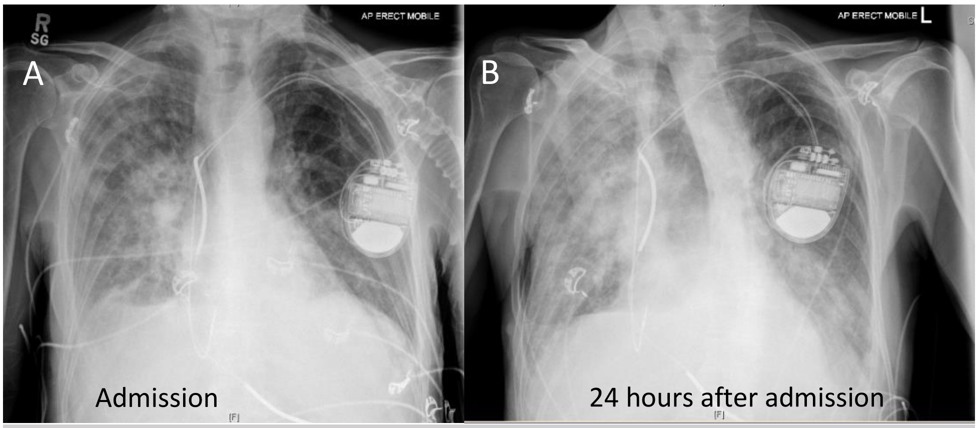


Figure 3. Chest X-rays of patient 6. Images at the time of admission and 24 hours later are shown as indicated.

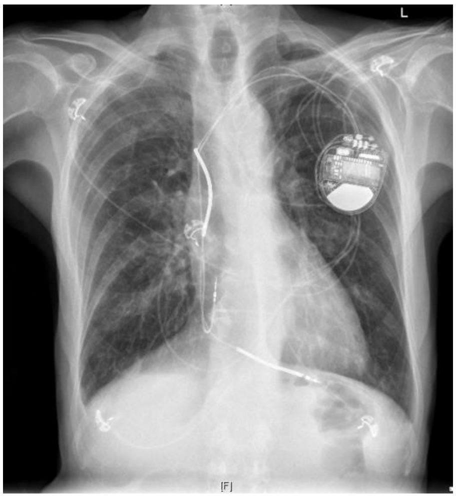

Figure 4. Chest X-ray of patient 6, taken 5 days after the episode of worsening pulmonary edema

At follow-up 6 weeks after discharge the patient had NYHA Class II symptoms, he had gained weight and unexplained increases in body temperature (that can be seen with low cardiac output) had not recurred. He was feeling well and had returned to work 2 weeks after discharge. The blood pressure was 100/80 mm Hg, the jugular venous pressure was not raised and there was no peripheral edema. The heart rate was 84 beats/min. The AM dose of carvedilol was increased to 6.25 mg while the nocte dose was maintained at 3.125 mg. Subsequently the carvedilol dose has been up-titrated 25 mg mane, 12.5 mg nocte. Further investigations of weight loss were cancelled. The patient has not been readmitted to hospital for 37 months for HF after mirabegron was introduced and functional NYHA Class II has been maintained.

***Patient 7***

An 80 year old male with a ~10 year history of idiopathic dilated cardiomyopathy developed worsening symptoms of HF despite maximal tolerated medical treatment. He had a LVEF of 20%. He had previously had an ICD. A recent attempt to upgrade to Bi-V pacing with a coronary sinus lead had been unsuccessful and he has admitted to hospital for insertion of a left ventricular epicardial lead.

Placement of the epicardial lead was uneventful but 3 days postoperatively he developed cardiogenic shock complicated by acute kidney injury and was admitted to the intensive care unit. Atrial flutter with rapid ventricular conduction was identified. He was treated with amiodarone initially given i.v. as well as with i.v. infusion of milrinone, norepinephrine and furosemide. Treatment also included vasopressin for a 24 hour period. He had successful DC cardioversion to sinus rhythm on day 4 postoperatively.

Acute kidney injury sustained during prolonged periods of hypotension gradually resolved and i.v. inotropic support was discontinued on day 6 postoperatively. The infusion of furosemide was replaced with oral administration, initially 40 mg t.d.s. The patient was transferred to a general cardiology ward on day 8 after placement of the pacing lead. His treatment for HF then consisted of carvedilol 12.5 mg b.d., furosemide 40 mg mane and spironolactone 12.5 mg b.d.

Signs of fluid overload and a deteriorating renal function with serum creatinine rising to 221 µmol/L re-developed over the next 5 days. The jugular venous pressure became raised to the angle of the jaw, the abdomen became distended with demonstrable shifting dullness attributed to ascites, and edema to the mid-thighs, sacral edema and basal pulmonary rales developed.

The patient was re-admitted to the intensive care unit on day 13 after Bi-V pacing was established and treatment with i.v. milrinone and i.v. furosemide was commenced. Treatment with levosimendan was considered but not used due to concern about the decrease in blood pressure it causes. Since severe HF had previously re-developed soon after inotropic support was weaned off treatment with mirabegron, 25 mg b.d. was prescribed 2 days after treatment with i.v. milrinone had been started. Blood pressure was maintained at 110/70 mm Hg on treatment with milrinone. An attempt at adding an ACE inhibitor to treatment was unsuccessful because it caused hypotension.

Diuresis and weight loss occurred with the treatment and clinical signs of fluid overload improved as did renal function. Milrinone was discontinued 4 days after readmission to the intensive care unit while treatment with mirabegron was maintained. The intensive care team had consulted the palliative care team for end-of-life discussion with the patient and his wife and it had been decided that no further i.v. inotropic support should be given if re-lapse of severe HF occurred.

Signs and symptoms of HF continued to improve after cessation of treatment with milrinone and the patient was transferred to the general ward after 6 days in the intensive care unit and discharged from hospital 4 days later. He had lost 10 kg in weight since readmission to the intensive care unit. Treatment for HF at discharge was carvedilol 12.5 mg b.d., furosemide 80 mg mane and spironolactone 12.5 mg b.d. and follow-up at home with a community-based palliative care team had been arranged.

On review 4 weeks after discharge the patient was ambulatory although limited by dyspnea on mild physical exertion. Physical signs of congestion had resolved. However, worsening symptoms developed and he was re-admitted to hospital with HF 10 months after treatment with mirabegron was started. After intensified conventional treatment he was again discharged but died from terminal HF 6 months later, 16 months after mirabegron had first been prescribed. Mirabegron had been discontinued 4 days before he died because renal function deteriorated to a level that precluded its use according to prescription guidelines.

***Patient 8***

A 68 year old male was admitted with worsening shortness of breath on minimal exertion. The jugular venous pressure was raised 5 cm above the sternal angle and moderate bilateral pitting edema was present to the knees. The BNP was elevated at 3150 ng/L. He had ischaemic cardiomyopathy and a history of myocardial infarction requiring emergency four-vessel coronary bypass surgery 11 years previously Echocardiography 1 year previously had estimated LVEF at 30%, and had shown mild aortic sclerosis and mild mitral -and tricuspid regurgitation. Treatment on admission included frusemide 40 mg mane, aspirin 150 mg mane, and carvedilol 12.5mg b.d. His blood pressure was 113/62 mm Hg. An ACE inhibitor had not been prescribed due to chronic renal impairment.

Repeated exacerbations of HF, of which this was the third precipitating admission in a year, occurred on a complex medical background. The patient had diabetes mellitus type 1 diagnosed aged 14 years. He had diabetic peripheral neuropathy with Charcot's deformity and amputation of 2 toes, vascular disease and diabetic retinopathy. He had also had a previous neurological transient ischemic attack. Chronic renal impairment was first detected 33 years previously and his baseline serum creatinine at the time of the index admission was 240 µmol/L. He had chronic lymphocytic leukaemia diagnosed for 12 years, recently complicated by white cell counts that had risen to ~70 X $10^9$/L.

The risk of compromising an already precarious renal function precluded aggressive treatment with furosemide. The usual dose of 40 mg mane was not increased and in view of the progressive HF with 3 recent admissions mirabegron, 25 mg mane was added. Signs and symptoms of HF improved to the point that treatment with furosemide was discontinued on request of his renal physician in the interest of optimising renal function, important in its own right but also important for allowing use of drugs to control of high white cell counts.

Discontinuation of furosemide was too ambitious and the patient presented again 9 days later with dyspnea. Furosemide was re-introduced at his usual daily dose of 40 mg and the dose of mirabegron increased with 25 mg nocte on alternate days in addition to the 25 mg mane dose. The patient again improved clinically and was discharged 4 days later. At discharge there was no elevation of jugular venous pressure, the chest was clear to auscultation and there were no signs of congestion on chest X-ray.

The patient has had admissions to hospital for treatment of foot ulcers and cellulitis and management of high white cell counts associated with his chronic lymphocytic leukemia was the dominant clinical issue until he was admitted with signs and symptoms of congestion 24 months after treatment with mirabegron was commenced. There was no clinically detectable precipitant for the cardiac de-compensation. However, he had a transient rise in serum troponin from a baseline of 18 µg/L (normal < 26 ng/L) to 804 ng/L indicating a clinically silent myocardial infarction. HF resolved and he was discharged with continued treatment with mirabegron, carvedilol 12.5 mg b.d. and furosemide 60 mg mane and 40 mg midday. Serum creatinine levels at the time of discharge were similar to those before mirabegron had been commenced 2 years previously. He remains in part-time work 32 months after mirabegron was first prescribed.

***Patient 9***

A 64 year old male with ischaemic cardiomyopathy presented with exacerbation of HF 1 month after a below knee amputation that had become necessary because of complications from diabetes mellitus type 2. The jugular venous pressure was raised to the jaw, and he had pitting edema to the groin and sacrum. He had gained 9 kg in weight and had shortness of breath and lethargy at rest. Echocardiography post-operatively showed a LVEF of 25%, moderate mitral regurgitation, a dilated right ventricle and pulmonary hypertension.

In addition to a 17 year history of diabetes mellitus, the patient had a history of myocardial infarction two years previously complicated by in-hospital ventricular fibrillation. An ICD was placed. He also had chronic renal impairment with baseline serum creatinine of 140 µmol/L.

Treatment initially included 80 mg i.v. furosemide but due to continued weight gain and an increase in serum creatinine a dobutamine infusion was added to treatment. This was gradually weaned off and eventually ceased 8 days after it had been started. After cessation of dobutamine the serum creatinine rose to 310 µmol/L over 4 days, edema to the sacrum re-developed and the jugular venous pressure became elevated to the angle of the jaw.

A decision was made to recommence treatment with dobutamine, start treatment with mirabegron once renal function had improved to a level at which it could be given according to prescription guidelines and only then wean off dobutamine while continuing treatment with mirabegron. Mirabegron 25 mg mane and 25 mg nocte on alternate days was commenced after 5 days' infusion of dobutamine. The blood pressure, on treatment with dobutamine just before treatment with mirabegron was commenced, was 121/75 mm Hg. Dobutamine was weaned off 3 days after treatment with mirabegron had been introduced.

After cessation of dobutamine infusion, the weight remained stable at 79 kg (from 93 kg at admission), peripheral pitting edema was markedly reduced. Serum creatinine levels were stable in a 230 -250 µmol/L range for a week after cessation of dobutamine. There was partial resolution of pleural effusions shown on chests X-rays but congestive changes remained (Figure 5). Clinically his HF had improved and he was discharged 12 days after treatment with mirabegron had been started.

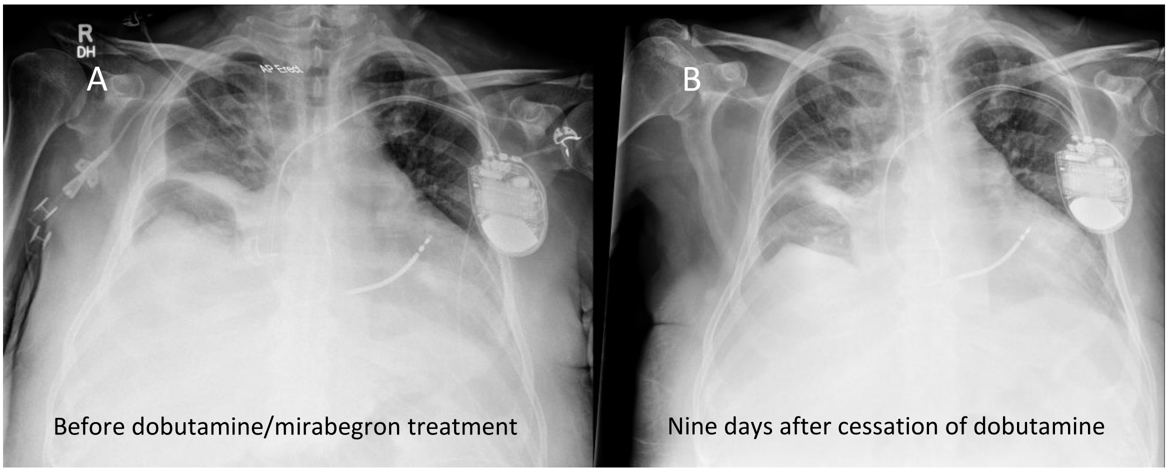


Figure 5. Chest X-rays of patient 9 after relapse of severe cardiac de-compensation with worsening acute-on-chronic renal impairment within 4 days after treatment with dobutamine showed pulmonary edema with a right pleural effusion with fluid tracking into the oblique fissure and a smaller sized left pleural effusion (panel A). The last available X-ray before discharge, 9 days after a second course of dobutamine treatment had been completed while treatment with mirabegron was maintained showed pleural effusions had been reduced in but not eliminated (panel B).

The patient was discharged to his home that was outside our Area Health Service and subsequent care was provided in hospitals other than ours. The patient's wife has reported to us that he continued to take the prescribed mirabegron and was reasonably well with no re-development of limb swelling for the first month after discharge. He was then admitted to a hospital for prosthetic limb rehabilitation. The limited information we have access to indicates that his serum creatinine during his admission there was 230 µmol/L, that is his renal function had been maintained at the level it was after his second treatment with dobutamine and treatment with mirabegron was commenced ~ 6 weeks previously at our hospital.

After discharge from the rehabilitation hospital the patient was later admitted to a university hospital outside our Area Health Service with a wound infection, worsening renal function and advanced HF. In contrast to what had been the case during his previous admission at our hospital volume overload and renal impairment was not responsive to i.v. inotropes and furosemide. He remained hypervolemic and developed anuric end-stage renal failure. Hemodialysis was started that presumably indicates his renal failure was not considered primarily due to HF. Mirabegron had been discontinued at that stage. He died from cardiac arrest during the admission, ~10 months after we had started treatment with mirabegron.

### SUPPLEMENTAL REFERENCES

1. William M, Vien J, Hamilton E, Garcia A, Bundgaard H, Clarke RJ and Rasmussen HH. The nitric oxide donor sodium nitroprusside stimulates the $Na^+$-$K^+$ pump in isolated rabbit cardiac myocytes. *J Physiol*. 2005;565:815-25.

2. Garcia A, Liu CC, Cornelius F, Clarke RJ and Rasmussen HH. Glutathionylation-Dependence of $Na^+$-$K^+$-Pump Currents Can Mimic Reduced Subsarcolemmal $Na^+$ Diffusion. *Biophys J*. 2016;110:1099-109.

3. Karimi Galougahi K, Liu CC, Garcia A, Fry NA, Hamilton EJ, Figtree GA and Rasmussen HH. beta3-Adrenoceptor activation relieves oxidative inhibition of the cardiac $Na^+$-$K^+$ pump in hyperglycemia induced by insulin receptor blockade. *Am J Physiol Cell Physiol*. 2015;309:C286-95.

4. Folland ED, Parisi AF, Moynihan PF, Jones DR, Feldman CL and Tow DE. Assessment of left ventricular ejection fraction and volumes by real-time, two-dimensional echocardiography. A comparison of cineangiographic and radionuclide techniques. *Circulation*. 1979;60:760-6.

5. Gadsby DC and Nakao M. Steady-state current-voltage relationship of the Na/K pump in guinea pig ventricular myocytes. *J Gen Physiol*. 1989;94:511-37.

6. Galougahi KK, Liu CC, Garcia A, Fry NA, Hamilton EJ, Rasmussen HH and Figtree GA. Protein kinase-dependent oxidative regulation of the cardiac $Na^+$-$K^+$ pump: evidence from in vivo and in vitro modulation of cell signalling. *J Physiol*. 2013;591:2999-3015.
